# Pathogenic KIF1A R350 Variants Disrupt A Conserved Kinesin-Tubulin Salt Bridge

**DOI:** 10.1101/2025.08.30.673262

**Authors:** Abhipsa Shatarupa, Lu Rao, Ana B. Asenjo, Arne Gennerich, Hernando Sosa

## Abstract

Pathogenic variants in the motor domain of the kinesin-3 motor protein KIF1A cause a range of neurodevelopmental and neurodegenerative conditions collectively termed KIF1A-associated neurological disorder (KAND). Among these, mutations at residue R350 are linked to hereditary spastic paraplegia and altered motor function. Yet, the structural basis for their pathogeny remains unclear. Here, we present high-resolution cryo-electron microscopy (cryo-EM) structures of KIF1A R350G and R350W bound to microtubules in both the apo and AMP-PNP-bound states. We identify a previously unrecognized salt bridge between KIF1A residue R350 and α-tubulin E415 that is disrupted in both mutants. This loss of electrostatic interaction correlates with increased velocity and reduced processivity, as demonstrated by single-molecule assays. Our results reveal a conserved electrostatic interaction at the motor–microtubule interface that regulates KIF1A’s motility behavior.

## Introduction

KIF1A is a neuron-specific, microtubule plus-end-directed motor protein critical for the anterograde transport of synaptic vesicle precursors and dense-core vesicles (*1-6*), as well as nuclear migration during brain development (*7, 8*). Mutations in the *KIF1A* gene are associated with a broad range of neurodevelopmental and neurodegenerative conditions, collectively referred to as KIF1A-associated neurological disorder (KAND) (*9, 10*).

Most KAND-associated mutations localize to the motor domain and are believed to impair motor function by disrupting ATP hydrolysis, microtubule binding, or the coordination between KIF1A’s two motor heads (*9, 11-16*). However, some mutations—such as V8M, A255V, and R350G— display gain-of-function features, including enhanced velocity or increased microtubule binding (*17, 18*), and have been identified in patients with hereditary spastic paraplegia (*19*).

To elucidate the structural and mechanistic consequences of mutations at residue R350, we determined cryo-EM structures of KIF1A R350G and R350W mutants in the apo and AMP-PNP-bound states and performed single-molecule fluorescence assays. Our findings reveal that both mutations disrupt a conserved salt bridge with α-tubulin, leading to increased velocity and reduced processivity.

## Results

### Cryo-EM structures of KIF1A R350 mutants bound to microtubules

We determine cryo-EM structures of dimeric KIF1A bearing KAND-associated mutations at R350 (R350G and R350W) in complex with microtubules, in both the nucleotide-free (apo) state and in the presence of the non-hydrolyzable ATP analog AMP-PNP (hereafter referred to as ANP) at resolutions ranging from 2.9 to 3.4 Å (Table 1, Supplementary Fig. 1).

**Table 1.** Cryo-EM Data Collection, Coordinate Refinement and validation statistics.

| **Dataset** | **MT-KIF1A-R350G-ANP** | **MT-KIF1A-R350W-ANP** | **MT-KIF1A-R350G-apo** | **MT-KIF1A-R350W-apo** |
| --- | --- | --- | --- | --- |
| PDB | TBD | TBD | TBD | TBD |
| EMDB | TBD | TBD | TBD | TBD |
| **Data collection and processing** |  |  |  |  |
| Microscope | EF-Krios | EF-Krios | EF-Krios | EF-Krios |
| Voltage (kV) | 300 | 300 | 300 | 300 |
| Camera | Gatan K3 | Gatan K3 | Gatan K3 | Gatan K3 |
| Magnification | 81000 | 64000 | 105000 | 105000 |
| Defocus range (um) | -0.6 to -2.5 | -0.8 to -2.5 | -0.5 to -2.3 | -0.1 to -2.3 |
| Exposure time (ms) | 1200 | 2000 | 1001 | 1320 |
| Electron dose (e-/Å<sup>2</sup>) | 50.27 | 52.14 | 52.88 | 57.96 |
| Calibrated pixel size (Å) | 0.8460 | 1.0730 | 0.8260 | 0.8260 |
| Revised pixel size (Å) of the final map | 0.8791 | - | - | - |
| Symmetry imposed | Helical | Helical | Helical | Helical |
| Rise (Å) | 5.453 | 5.536 | 5.562 | 5.571 |
| Twist (deg) | 168.07 | 168.07 | 168.07 | 168.07 |
| Initial particle images after symmetry expansion (no.) | 5,286,915 | 4,391,445 | 4,318,200 | 2,376,645 |
| Final particle images (no.) | 1,194,267 | 163,145 | 4,318,200 | 953,398 |
| Overall Map resolution (Å), 0.143 FSC | 2.95 | 3.30 | 3.29 | 3.21 |
| Map sharpening B-factor (Å<sup>2</sup>) | 84.3 | 62.4 | 114.1 | 104.8 |
| **Refinement** |  |  |  |  |
| **Model composition** |  |  |  |  |
| Non-hydrogen atoms | 20146 | 20166 | 9741 | 9751 |
| Protein residues | 2530 | 2530 | 1227 | 1227 |
| Ligands | 11 | 11 | 4 | 4 |
| **B factors (Å<sup>2</sup>)** |  |  |  |  |
| Protein | 121.44 | 127.74 | 117.38 | 113.59 |
| Ligand | 100.89 | 133.02 | 118.70 | 115.91 |
| **RMS deviations** |  |  |  |  |
| Bond lengths (Å) | 0.006 | 0.002 | 0.005 | 0.005 |
| Bond angles (°) | 1.181 | 0.504 | 0.831 | 1.121 |
| **Validation** |  |  |  |  |
| MolProbity score | 1.28 | 1.70 | 1.24 | 1.12 |
| Clash score | 1.39 | 6.68 | 2.87 | 1.09 |
| Rotamer outliers (%) | 1.43 | 1.47 | 0.47 | 0.85 |
| **Ramachandran plot** |  |  |  |  |
| Favored (%) | 95.79 | 96.74 | 97.05 | 95.58 |
| Allowed (%) | 4.17 | 3.18 | 2.87 | 4.42 |
| Outliers (%) | 0.04 | 0.08 | 0.08 | 0.00 |

In the ANP-bound state, both R350 mutants adopt a two-heads-bound configuration resembling that of wild-type KIF1A (*15*) (Fig. 1a,b). The leading head assumes an open conformation with a rear-pointing (minus-end-directed) neck-linker, while the trailing head is in a closed conformation with a docked neck-linker. Both heads show clear nucleotide density in their active sites. Moreover, in these structures, densities for the α- and/or β-tubulin C-terminal tails interacting with the motor domain’s K-loop are well resolved, indicating that the R350 mutations do not perturb the K-loop-tubulin interaction.

**Fig. 1.**
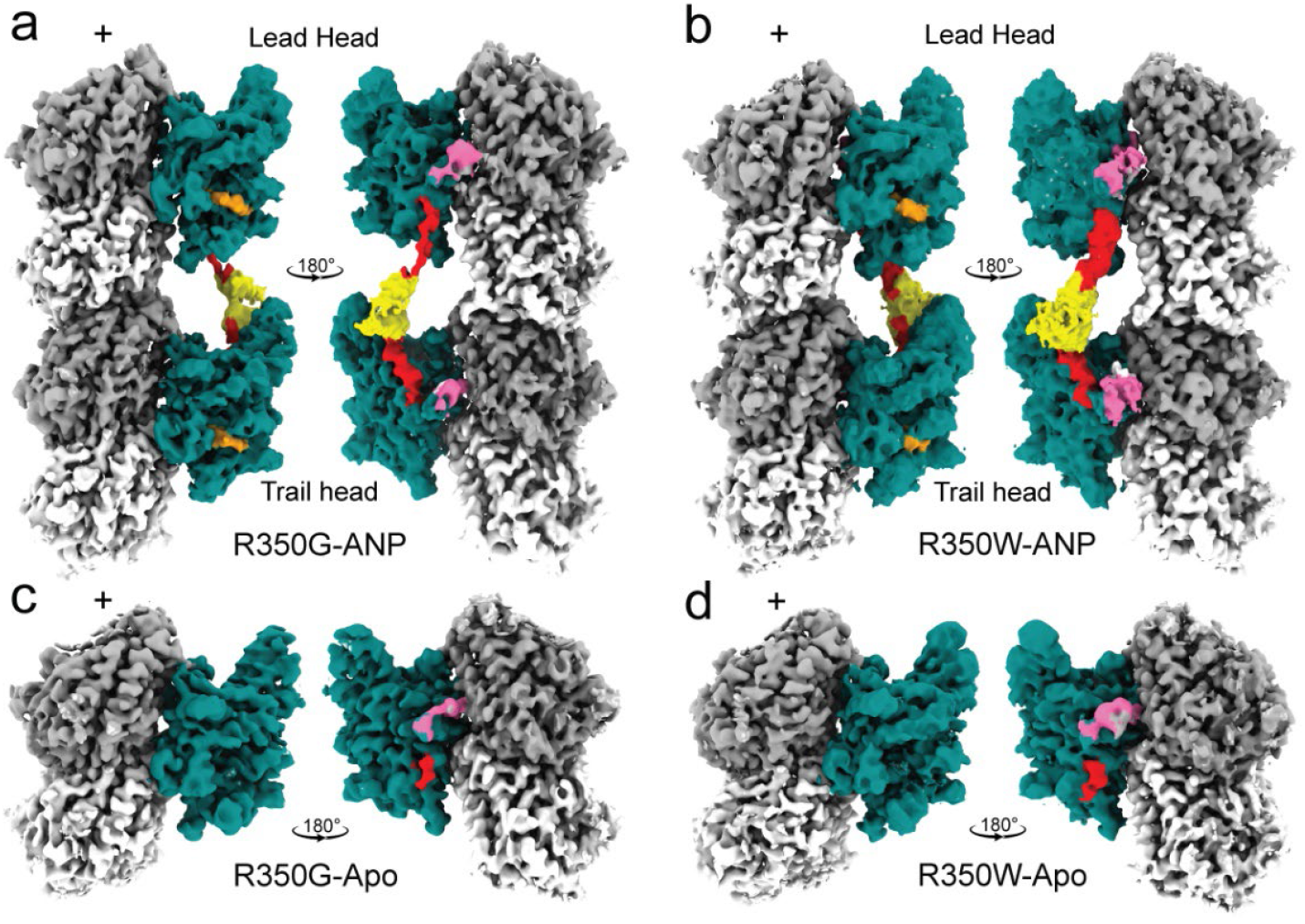
**a-d**. Isosurface representations of 3D maps for microtubule-bound KIF1A-R350G and KIF1A-R350W mutants in the ANP and apo states. Surface coloring highlights distinct structural elements: KIF1A core motor domain in teal, K-loop (loop-12) in pink, neck-linker in red, coiled-coil neck in yellow, and ANP in orange, α-tubulin in light gray, and β-tubulin in dark gray. Densities corresponding to the K-loop were low-passed filtered and displayed at a lower contour level to better visualize this flexible region. Densities for the coiled-coil and parts of neck-linker regions are absent in the apo state (c-d).

In the apo state, both mutants adopt a one-head-bound configuration with the bound motor domain in an open conformation like apo wild-type KIF1A (*15*) (Fig. 1c,d). Only the initial segment of the minus-end-directed neck-linker is visible, while the remainder of the neck-linker, the neck coiled-coil, and the partner motor domain are not resolved, indicating a high flexibility in the unbound partner motor head.

### R350 mutations selectively disrupt a salt bridge in the open conformation

The open and closed conformations of the nucleotide-binding pocket can be characterized by measuring the average distances between highly conserved kinesin residues involved in nucleotide coordination (*15, 20*). These distances in both conformational states were very similar between wild type KIF1A and the R350G and R350W mutants (Fig. 2a), indicating that overall nucleotide-binding geometries remain intact.

**Fig. 2.**
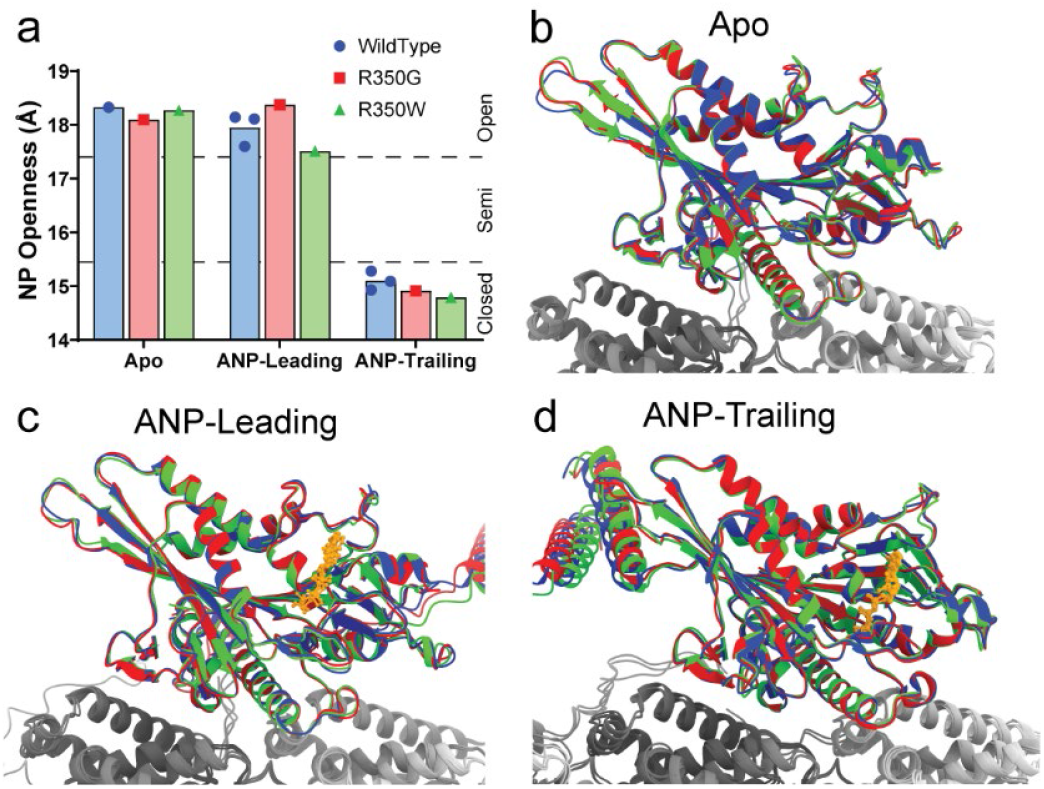
KIF1A motor domain structures of wild type and R350 mutants. **a**. Average distance between highly conserved residues across the nucleotide-binding pocket (R216 and A250 to P14, S104, and Y105). **b–c**, Structural alignment of KIF1A WT (blue), R350G (red), and R350W (green) motor domains in the apo state (b), the leading head of the ANP state (c), and the trailing head of the ANP state (d).

Structural comparison of wild type and mutant motor domains shows that, aside from the altered residue at position 350, the structures are nearly superimposable in their respective conformations, including regions surrounding the mutation site (Fig. 2b-d). The most prominent structural difference lies in the chemical nature of residue 350 (Fig. 3). In the open conformation, the wild-type R350 resides near α-tubulin residue E415 at a distance compatible with the formation of a salt bridge (Fig. 3a,b). However, in the closed conformation, R350 is further away from α-tubulin E415 and does not contact tubulin (Fig. 3c).

**Fig. 3.**
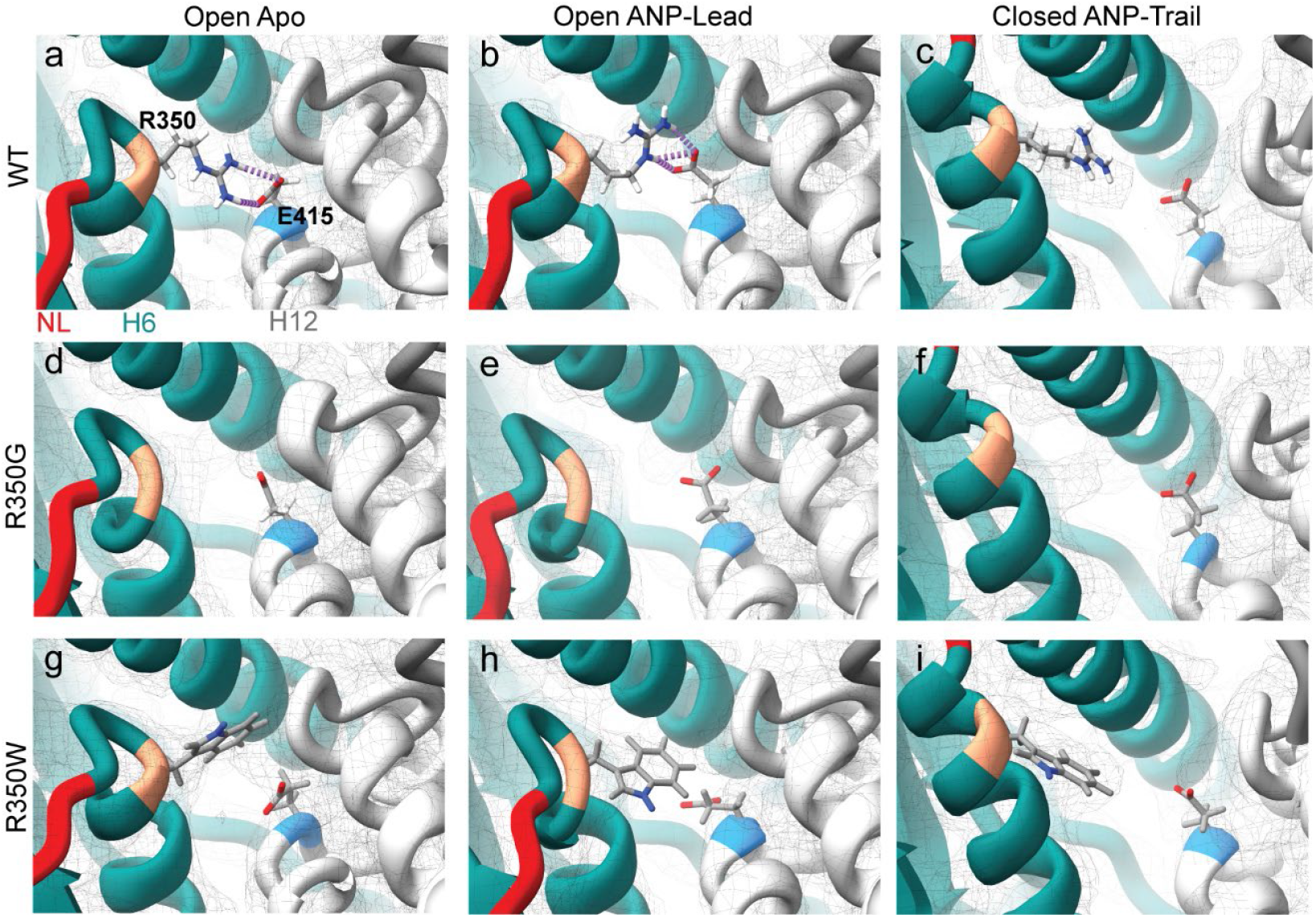
Structures of microtubule-bound KIF1A WT and R350 mutants near residue 350. **a-c**. WT. **d-f**. R350G. **g-i**. R350W. Apo state, open conformation (a, d, g); ANP state, leading head open conformation (b, e, h); ANP state, trailing head closed conformation (c, f, i). Structures are shown in ribbon representation with the KIF1A motor domain in teal, the neck linker in red, and residue 350 in salmon. α-tubulin is shown in light gray, with residue E415 in blue. Side chains of KIF1A residue 350 and α-tubulin E415 are shown as sticks and colored by heteroatom. Hydrogen bonds between KIF1A WT R350 and α-tubulin E415 are indicated as dashed purple lines. Corresponding cryo-EM densities are shown as semitransparent grey mesh isosurfaces. WT structures correspond to PDB IDs 8UTS (a) and 8UTN (b–c) after molecular dynamics fitting with ISOLDE (*38*).

In the R350G mutant, the side chain is absent, and in the R350W mutant, the positively charged arginine is replaced with a bulky hydrophobic residue. Both substitutions eliminate the open conformation salt bridge (Fig. 3d,e,g,h). These findings identify a previously unrecognized interaction between the motor domain and α-tubulin that is conformation-dependent and likely relevant to motor function.

### R350 mutations increase velocity and reduce processivity

To assess the functional consequences of the R350 mutations, we performed single-molecule motility assays to measure two key parameters of kinesin movement: velocity and run length (Fig. 4). Velocity reflects the rate of forward stepping, while run length is the average distance traveled before detachment and serves as a proxy for processivity, the motor’s ability to take multiple successive steps without dissociating.

**Fig. 4.**
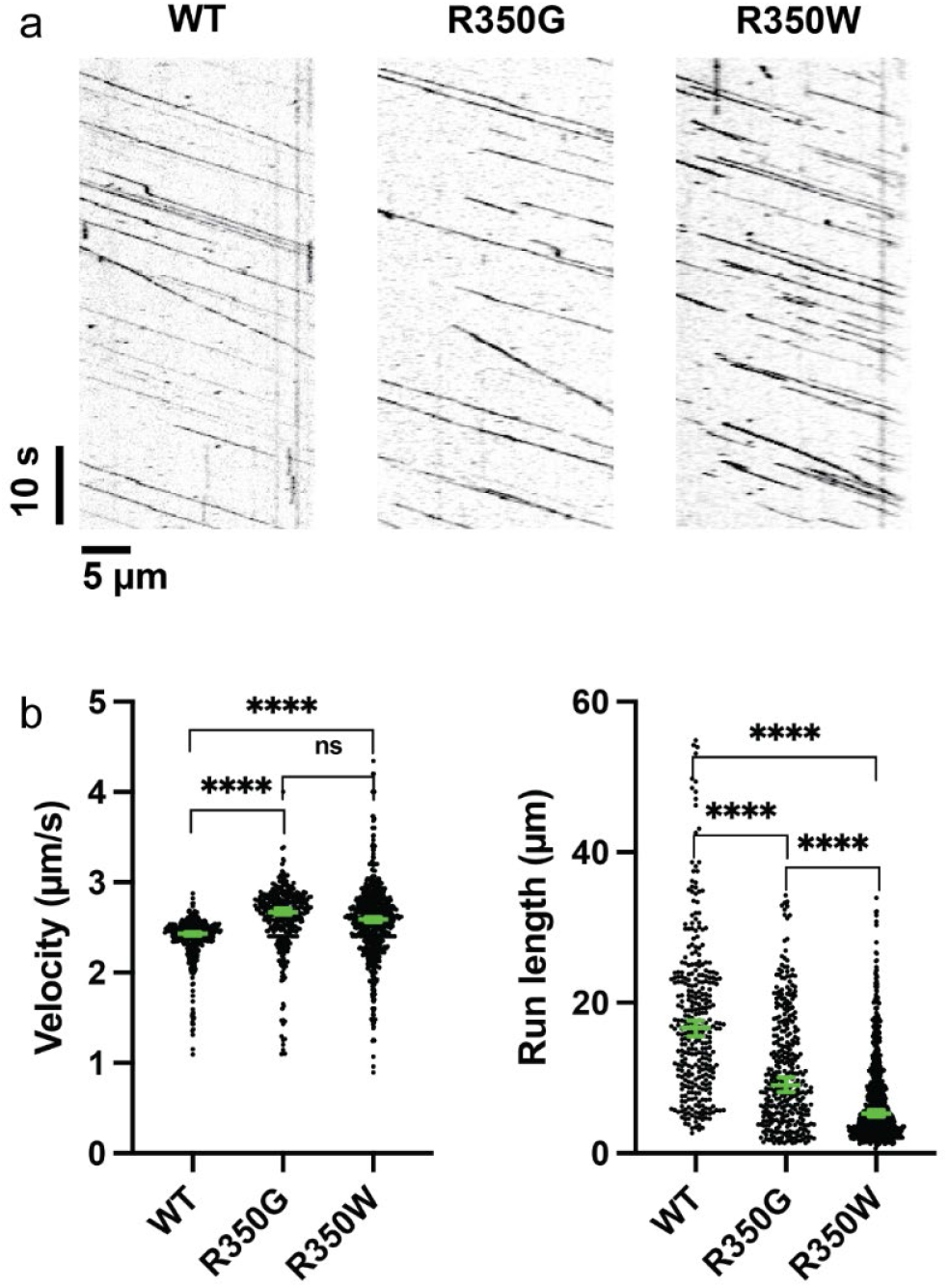
Single-molecule behaviors of KIF1A WT and R350 mutants. **a**. Examples of kymographs of KIF1A WT, R350G, and R350W. **b**. Left panel: velocity of KIF1A WT, R350G, and R350W. The green bars indicate median value with 95% confidence interval (CI). WT: 2.43 [2.41, 2.45] μm/s, n=330. R350G: 2.67 [2.64, 2.71] μm/s, n=354. R350W: 2.59 [2.56, 2.62] μm/s, n=673. Welch’s t-test was performed for the constructs. Two-tailed P value: ^****^, P<0.0001; ns, P=0.0774. Right panel: Processivity of KIF1A WT, R350G, and R350W. The green bars indicate median value with 95% confidence interval (CI). The numbers of data points are the same as the velocity plot. WT: 16.7 [15.5, 17.6] μm. R350G: 9.0 [8.1, 10.0] μm. R350W: 5.3 [4.9 5.7] μm. Kolmogorov-Smirnov test was performed for the constructs. ^****^, P<0.0001.

Both R350G and R350W mutants exhibited significantly increased velocities compared to wild-type KIF1A (Fig. 4b), consistent with previous reports showing that R350G enhances stepping speed and can rescue locomotion defects in *C. elegans* lacking the corresponding endogenous kinesin (*17*). In contrast, the run lengths were significantly reduced for both mutants (Fig. 4b), indicating impaired processivity. The reduction was more pronounced for R350W. Structural comparison of both mutants in the presence of ANP reveals subtle differences in the trajectory of the backward-oriented neck linker in the leading head and in the orientation of the trailing motor domain (Fig. 5). We hypothesize that these structural differences alter inter-head tension and reduce coordination between the motor domains, contributing to the more pronounced decrease in run lengths observed for the R350W mutant.

**Fig. 5.**
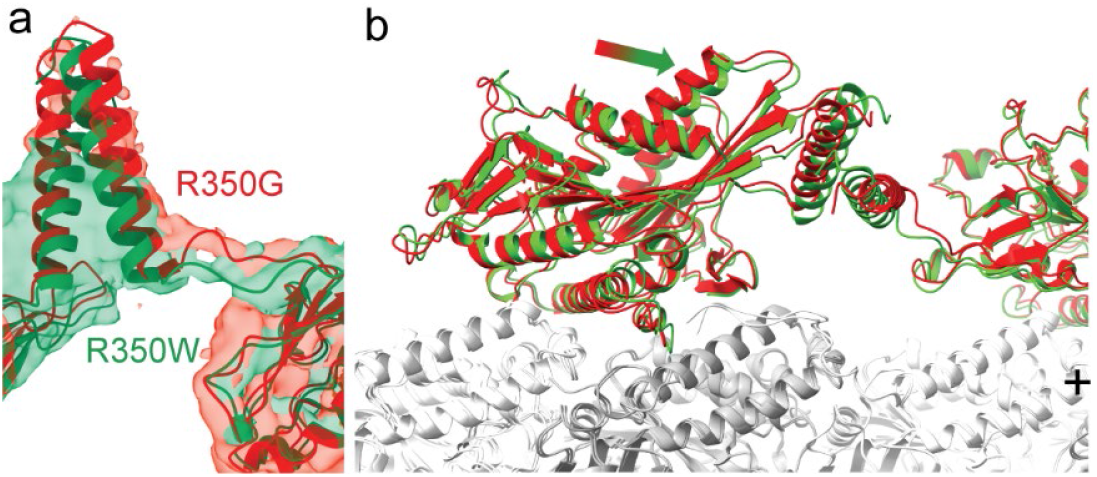
Comparison of KIF1A R350G and R350W mutants. **a**. ANP state leading-head neck-linker path of R350G (red) and R350W (green). Structures are shown in cartoon representation with corresponding cryo-EM densities displayed as semitransparent isosurfaces. Both mutant structures are aligned to the leading-head motor domain. **b**. Trailing-head conformations of R350G (red) and R350W (green) bound to microtubules. Structures are shown in cartoon representation, with α-tubulin in light gray and β-tubulin in dark gray. Both structures are aligned to the corresponding tubulin subunits. The arrow indicates that R350W is positioned slightly more forward (towards the microtubule plus end) relative to R350G.

Together, these results suggest that disruption of the KIF1A R350 to α-tubulin E415 salt bridge lowers the energetic barrier for the open-to-closed transition, increasing stepping rate. Conversely, the absence of this salt bridge may increase the likelihood of leading head (open conformation) detachment and, consequently, the probability of both heads detaching simultaneously during translocation, resulting in reduced processivity (Fig. 6).

**Fig. 6.**
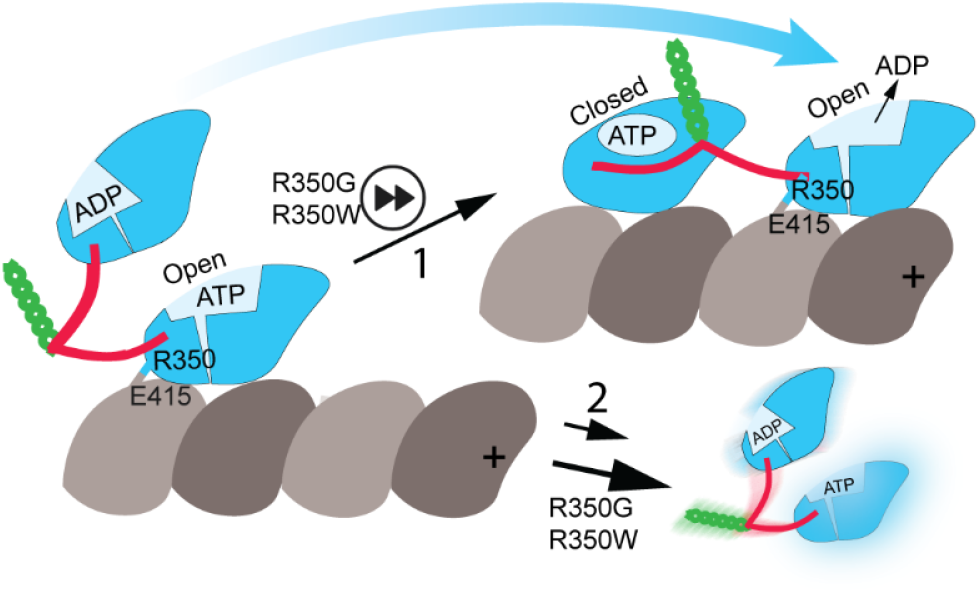
Model of altered motor function in KIF1A R350 mutants. KIF1A residue R350 forms a salt bridge with α-tubulin E415 in the open conformation of the motor domain. Disruption of this salt bridge in the R350G and R350W mutants facilitates conversion to the closed state, with neck-linker docking and forward movement of the partner motor domain to the leading position (step 1), resulting in increased translocation velocity. Conversely, disruption of the salt bridge also increases the likelihood of microtubule detachment (step 2), leading to reduced processivity. KIF1A motor domain is shown in blue, neck-linker in red, neck coiled coil in green, α-tubulin in light grey, and β-tubulin in dark grey. The microtubule plus end is oriented toward the right.

## Discussion

Here, we solved the cryo-EM structures of dimeric, microtubule-bound KIF1A carrying disease-associated mutations at position R350 (R350G and R350W) and characterized their motility using single-molecule assays. We found that these mutations cause modest yet significant alterations in motor behavior, specifically, increased velocity and decreased processivity, due to the disruption of a previously unrecognized electrostatic interaction between KIF1A R350 and α-tubulin residue E415.

The R350G mutation has been reported to produce a relatively mild clinical phenotype compared to other KAND-associated mutations (*9*), which is consistent with the subtle structural and functional changes we observed. There is less data regarding the R350W mutation, but it has been reported as a dominant mutation (*21*), suggesting that even moderate motility defects can manifest as disease in the heterozygous state.

We propose that the primary pathogenic mechanism of R350 mutations arises from the loss of a stabilizing salt bridge formed in the open, microtubule-bound conformation of KIF1A between R350 and α-tubulin E415. The resulting increase in motor stepping rate and decrease in run length suggest a reduced energetic barrier for the open-to-closed transition and an elevated likelihood of detachment during stepping. KIF1A residue R350 is located near the end of α-helix 6, at a position where a positively charged residue (arginine or lysine) is highly conserved across the kinesin superfamily. Although the role of this residue in kinesin–microtubule interactions has not previously been examined, its conservation suggests that it likely plays a similar role in other kinesins. Notably, a recent study reported that the complementary mutation in α-tubulin 4A, E415K, results in spastic ataxia (*22*). These observations emphasize the critical role of this electrostatic interaction in kinesin motility and its potential contribution to disease pathology.

An earlier hypothesis proposed that the R350G mutation alters KIF1A activity by affecting the neck-linker conformation, based on the proximity of R350 to the neck-linker (*19, 23*). However, this idea predated high-resolution (<4 Å) kinesin–microtubule structures (*15, 20*) and assumed that KIF1A R350 does not interact with the microtubule, which is not the case, as highlighted in this work. Furthermore, our data show nearly identical neck-linker conformations between wild-type and the R350G mutant in both the open and closed states, indicating no substantial structural effect of this mutation on the neck-linker.

Another possibility is that the R350G mutation activates the motor by disrupting the autoinhibited, folded conformation of full-length KIF1A, as has been proposed for other gain-of-function mutations like V8M and A255V (17). While we cannot completely rule out this mechanism, structural considerations argue against it. A proposed structural model for kinesin-3 autoinhibition is based on the crystal structure of full-length KLP6 (*24*). This structure shows a compacted folded conformation of the kinesin monomer where the coiled coils and FHA (forkhead-associated domain) domains fold onto the motor domain, covering the nucleotide-binding site and preventing dimerization. In this structure, the residue equivalent to KIF1A R350 (R354 in KLP6) is located far from the regions involved in forming the folded autoinhibited state and does not contact any structural elements associated with autoinhibition (Supplementary Fig. 2a). Similarly, predicted full-length models of KIF1A using Aphafold3 (*25*) place R350 far from the key intramolecular interfaces involved in forming the autoinhibited conformation (Supplementary Fig. 2b,d,e), further supporting the conclusion that R350 is unlikely to contribute to autoinhibition. It has also been proposed that KIF1A in the autoinhibited state exists as a dimer rather than a monomer (*26*), but AlphaFold3 dimer models likewise place residue R350 far from the interfaces that form the folded autoinhibited state (Supplementary Fig. 2c,f,g). Taken together, these structural considerations support our conclusion that the disruption of the open-conformation KIF1A R350 to α-tubulin E415 salt bridge is the primary mechanism underlying the altered motor function of KIF1A R350 mutations.

In summary, our findings demonstrate that mutations at R350 perturb a critical salt-bridge at the motor-microtubule interface, leading to enhanced velocity and reduced processivity. The fact that such subtle biophysical changes result in clinically observable motor dysfunction underscores the importance of precise kinetic and mechanical tuning in neuronal transport. These insights not only clarify the molecular basis of KAND-related R350 mutations but also suggest broader relevance for understanding how microtubule-motor interface integrity contributes to neuronal health.

## Methods

### Generation of plasmids for KIF1A constructs

A plasmid from a previous published KIF1A construct (*9*) (KIF1A (*Homo sapiens*, aa 1-393)-leucine zipper-SNAPf-EGFP-6His) was used as the template for all constructs in this study. For proteins used in the cryo-EM studies, the SNAPf-EGFP-6His tag was replaced with a strep-II tag (IBA Lifesciences GmbH) using Q5 mutagenesis (New England Biolabs Inc., #E0554S) as in a previous study(*15*). Mutations within KIF1A were generated using Q5 mutagenesis. All plasmids were confirmed by Sanger sequencing (Albert Einstein College of Medicine, Genomic Core Facility).

### Protein expression in *E. coli*

KIF1A expression was performed as follows: Each plasmid was transformed into BL21-CodonPlus(DE3)-RIPL competent cells (Agilent Technologies, #230280). A single colony was picked and inoculated in 1 mL of terrific broth (TB) (protocol adopted from Cold Spring Harbor protocol (*27*) with 50 µg/mL carbenicillin and 50 µg/mL chloramphenicol. The 1-mL culture was shaken at 37 °C overnight and then inoculated into 400 mL of TB (or 1–2 L for cryo-EM studies) with 2 µg/mL carbenicillin and 2 µg/mL chloramphenicol. The culture was shaken at 37 °C for 5 hours and then cooled on ice for 1 hour. Afterwards, IPTG was added to the culture to a final concentration of 0.1 mM and the culture was shaken at 16 °C overnight to induce protein expression. The cells were harvested by centrifugation at 3,000×g for 10 minutes at 4°C. The supernatant was discarded, and 1.25 mL of B-PER™ Complete Bacterial Protein Extraction Reagent (ThermoFisher Scientific, #89821) per 100 mL culture with 2 mM MgCl_2_, 1 mM EGTA, 1 mM DTT, 0.1 mM ATP, and 2 mM PMSF was added to the cell pellet. The cells were fully resuspended, and flash frozen in liquid nitrogen. If the purification was not done on the same day, the frozen cells were stored at –80 °C.

### Protein purification

To purify the protein, the frozen cell pellet was thawed at 37 °C. The solution was nutated at room temperature for 20 minutes and then dounced for 10 strokes on ice to lyse the cells. Unless specified, the following procedures were done at 4 °C. The cell lysate was cleared by centrifugation at 80,000 rpm (260,000 ×g, *k*-factor=28) for 10 minutes in a TLA-110 rotor using a Beckman Tabletop Ultracentrifuge Unit. The supernatant was flown through 500 μL of Roche cOmplete™ His-Tag purification resin (Millipore Sigma, #5893682001) for His-tag tagged proteins, or 2 mL of Strep-Tactin® 4Flow® high capacity resin (IBA Lifesciences GmbH, #2-1250-010) for strep-II tagged proteins. The resin was washed with wash buffer (WB) (for His-tagged protein: 50 mM HEPES, 300 mM KCl, 2 mM MgCl_2_, 1 mM EGTA, 1 mM DTT, 1 mM PMSF, 0.1 mM ATP, 0.1% Pluronic F-127 (w/v), 10% glycerol, pH 7.2; for strep-II tagged protein, Pluronic F-127 and glycerol were omitted). For proteins with a SNAPf-tag, the resin was mixed with 10 μM SNAP-Cell® TMR-Star (New England Biolabs Inc., #S9105S) at room temperature for 10 minutes to label the SNAPf-tag. The resin was further washed with WB, and then eluted with elution buffer (EB) (for His-tagged protein: 50 mM HEPES, 150 mM KCl, 150 mM imidazole, 2 mM MgCl_2_, 1 mM EGTA, 1 mM DTT, 1 mM PMSF, 0.1 mM ATP, 0.1% Pluronic F-127 (w/v), 10% glycerol, pH 7.2; for strep-II tagged protein: 80 mM PIPES, 2 mM MgCl_2_, 1 mM EGTA, 1 mM DTT, 0.1 mM ATP, 5 mM desthiobiotin). The Ni-NTA elute was flash frozen and stored at –80 °C. The Strep-Tactin elute was concentrated using a Amicon Ultra-0.5 mL Centrifugal Filter Unit (30-kDa MWCO) (Millipore Sigma, #UFC503024). Storage buffer (SB) (80 mM PIPES, 2 mM MgCl_2_, 1 mM EGTA, 80% sucrose (w/v)) was added to the protein solution to have a final 20% sucrose (w/v) concentration, and the protein solution was flash frozen and stored at -80 °C. The purity of the proteins was confirmed on polyacrylamide gels (Supplementary Fig. 3).

### Microtubule-binding and -release assay

A microtubule (MT)-binding and -release (MTBR) assay was performed to remove inactive motors for single-molecule TIRF assay. 50 μL of eluted protein was buffer-exchanged into a low salt buffer (30 mM HEPES, 50 mM KCl, 2 mM MgCl2, 1 mM EGTA, 1 mM DTT, 1 mM AMP-PNP, 10 µM taxol, 0.1% Pluronic F-127 (w/v), and 10% glycerol) using 0.5-mL Zeba™ spin desalting column (7-kDa MWCO) (ThermoFisher Scientific, #89882). The solution was warmed to room temperature and 5 μL of 5 mg/mL taxol-stabilized MTs were added. The solution was well mixed and incubated at room temperature for 2 minutes to allow motors to bind to the MTs. Afterward, the solution was spun through a 100 μL glycerol cushion (80 mM PIPES, 2 mM MgCl_2_, 1 mM EGTA, 1 mM DTT, 10 µM taxol, and 60% glycerol, pH 6.8) by centrifugation at 45,000 rpm (80,000×g, *k*-factor=33) for 10 minutes at room temperature in TLA-100 rotor using a Beckman Tabletop Ultracentrifuge Unit. Next, the supernatant was removed, and the pellet was resuspended in 50 μL high salt release buffer (30 mM HEPES, 300 mM KCl, 2 mM MgCl_2_, 1 mM EGTA, 1 mM DTT, 10 μM taxol, 3 mM ATP, 0.1% Pluronic F-127 (w/v), and 10% glycerol). The MTs were then removed by centrifugation at 40,000 rpm (60,000×g, *k*-factor=41) for 5 minutes at room temperature. Finally, the supernatant containing the active motors was aliquoted, flash frozen in liquid nitrogen, and stored at –80 °C.

### Single-molecule TIRF motility assay

MTBR fractions were used for the single-molecule TIRF assay, and the dilutions were adjusted to an appropriate density of motors on MTs. The assay was performed as follows: A flow chamber was assembled with a glass slide (Fisher #12-550-123), an ethanol-cleaned coverslip (Zeiss #474030-9000-000), and two stripes of parafilm. All the following incubation was done at room temperature. 10 µl of 0.5 mg/ml BSA-biotin was flown into the chamber with 10 min incubation. The chamber was then washed with 2×20 µl blocking buffer (80 mM PIPES, 2 mM MgCl_2_, 1 mM EGTA, 10 µM taxol, 1% Pluronic F-127 (w/v), pH 6.8) and incubated for 10 min to block the surface. Afterwards, 10 µl of 0.25 mg/ml streptavidin was introduced into the chamber with 10 min incubation. The chamber was washed with 2×20 µl blocking buffer, and 10 µl of 0.02 mg/ml Cy5- and biotin-labeled MTs was flown into the chamber with 1 min incubation. The chamber was washed with 2×20 µl blocking buffer. The MTBR motor was diluted in motility buffer (80 mM PIPES, 2 mM MgCl_2_, 1 mM EGTA, 1 mM DTT, 10 µM taxol, 0.5% Pluronic F-127 (w/v), 2 mM ATP, 5 mg/mL BSA, 1 mg/mL α-casein, gloxy oxygen scavenging system, and 10% glycerol, pH 6.8) in an appropriated dilution, and the solution was introduced into the chamber. The chamber was sealed with vacuum grease. Images were acquired with 200 ms per frame (total 600 frames per movie) and then analyzed via a custom-written MATLAB software. Kymographs were generated using ImageJ2 (version 2.16.0). GraphPad Prism (version 10.4.2) was used to perform statistical analysis and generate graphs.

### Cryo-EM

#### Preparation of microtubules

Microtubules (MTs) were prepared from porcine brain tubulin (Cytoskeleton, Inc. CO) as described in (*15*).

#### Preparation of MT-KIF1A complexes

Four µL of ∼6 μM MT solution in BRB80 buffer (80 mM PIPES, 2 mM MgCl_2_, 1 mM EGTA, Ph 6.8) plus 20 μM paclitaxel were layered onto (UltrAuFoil R1.2/1.3 300 mesh) plasma cleaned grids. The MTs were incubated for 1 minute at room temperature and then the excess liquid removed from the grid using Whatman #1 paper. Four µL of a solution containing either KIF1A-R350G (20 μM) or KIF1A-R350W (17.5 μM) in BRB80 supplemented with 20 μM paclitaxel and either 5 mM AMP-PNP (ANP conditions) or 5×10^-3^ units per μL apyrase (Apo conditions). The grid with the MT and kinesin mixture were then mounted into a Vitrobot apparatus (FEI-ThermoFisher MA), incubated 1 min at room temperature and plunge-frozen into liquid ethane (Vitrobot settings: 100% humidity, 3 seconds blotting with Whatman #1 paper and -2 mm offset). Grids were then transferred and stored into liquid nitrogen until data collection.

#### Cryo-EM data collection

All data were collected on a TF Titan Krios operating at 300 kV and equipped with a Gatan K3 Summit direct electron detector (Gatan, Pleasanton, CA). Data collection was controlled using Leginon (*28, 29*). The pixel size, defocus range, and total accumulated dose are summarized in Table 1.

#### Cryo-EM Image analysis and 3D reconstruction

All cryo-EM datasets were processed in CryoSPARC v.4.6.0 (*30*) using an adapted version of the helical single-particle reconstruction and helical assembly subunit refinement/classification (HASRC) workflow (*31*). Initial processing for each data set involved patch motion correction and contrast transfer function (CTF) estimation and correction. After micrographs curation, microtubules were manually picked, followed by 2D classification and use of 2D templates to select filament particles using filament tracer. Filament particles were extracted with filament diameter of 350 Å and box size of 664 x 664 pixels. 2D classification of all the filament particles were performed. At this step, class averages corresponding to different microtubule types were distinguished based on their diameter and Moiré pattern (32), and particles corresponding to 15-protofilament microtubules with a right-handed twist (15R) were selected for further processing. Helical refinement was then done with the selected particles using as initial values a helical twist of 168.07°, a helical rise of 5.6 Å, symmetry order of 15 and a maximum out-of-plane tilt of 15° (final twist and rise refined values listed in Table 1). Symmetry expansion was performed using the output parameters from the helical refinement and 15 symmetry order. To reduce computation time, after symmetry expansion the larger datasets were randomly subdivided, and subsets of 4.5 million particles or fewer were selected for further processing. The symmetry expanded dataset were then subjected to signal subtraction to isolate individual protofilaments with associated kinesins. For these, masks encompassing at least 3 tubulin heterodimers and associated kinesins were created. The signal-subtracted single-protofilament reconstruction was locally refined using a mask that encompassed only the tubulin region of the selected protofilament. The resulting 3D reconstruction was then centered and cropped around the isolated protofilament. This dataset was then subsequently subjected to 3D classification to separate classes with and without kinesin decoration, as well as distinct motor domain conformations. For 3D classification, the data were first downsampled to a pixel size of ∼7.5 Å/pixel using Fourier cropping.

For the ANP datasets, where two-head microtubule-bound states were expected, two rounds of classification into six classes were performed. The first round used a mask positioned at the leading site of the protofilament, and the second round used a mask at the trailing site. Both masks encompassed the volume of a motor domain together with the coiled-coil region, modeled in either the “up” position (at the plus end of the mask volume) or the “down” position (at the minus end of the mask volume). Each classification yielded classes corresponding to: (i) leading-like configurations, with coiled-coil densities in the down position; (ii) trailing-like configurations, with coiled-coil densities in the up position; (iii) empty classes with no discernible kinesin densities; and (iv) noisy classes that could not be confidently assigned as leading, trailing, or empty. Particles classified as leading at the leading position and trailing at the trailing position were selected using the Particle Sets tool. A 3D reconstruction of this data subset was generated and locally refined using a mask encompassing two tubulin heterodimers and the bound kinesins. For the apo dataset, only one round of 3D classification was performed on a single kinesin motor domain, since the helical reconstruction already indicated that these datasets corresponded to kinesins with only one microtubule-bound head. The classification therefore yielded classes showing either bound kinesin densities or empty sites, with the bound kinesins showing no coiled-coil densities in either the up or down position. Masks for local refinement, 3D classification, and inverse masks for particle subtraction were generated from low-resolution (30 Å) density maps based on atomic models of kinesin-decorated protofilaments. These were produced using the molmap function in UCSF Chimera (*32*), and then imported, thresholded, and inverted (for particle subtraction) using the CryoSPARC volume import and volume tools.

#### Model building and model refinement

Initial models PDB: 8UTN and PDB: 8UTS (*15*) were fitted into the ANP and apo maps, respectively, using the *Fit in Map* function in UCSF Chimera (*33*). Mutations at residue R350 were introduced in Coot (*34*), replacing it with glycine (R350G) or tryptophan (R350W). These initial mutated models for each of the four complexes were flexibly fitted into their respective cryo-EM maps using Rosetta refinement protocols (*35*). Models with the best agreement to the cryo-EM density and best Molprobity (*36*) scores were selected and further refined using Phenix real-space refinement (*37*). Refined models were then manually edited using Coot (*34*) and ISOLDE (*38*). Several rounds of manual editing and Phenix real-space refinement were performed until the final models were obtained. Figures of the atomic models and cryo-EM maps were prepared using ChimeraX (*32*).

## Acknowledgements

This work was supported by National Institutes of Health Grant R01GM147332 (A.G and H.S), R01GM113164 (H.S.). Cryo-EM data collection was performed at the Simons Electron Microscopy Center and National Resource for Automated Molecular Microscopy located at the New York Structural Biology Center, supported by grants from the Simons Foundation (SF349247), NYSTAR, and the NIH National Institute of General Medical Sciences (GM103310) with additional support from Agouron Institute (F00316) and NIH (OD019994). We thank Wendy Chung for helpful discussions on disease severity of KAND-associated R350 mutations.

## Data availability

The data that support this study are available from the corresponding authors upon request. The structural models generated in this study have been deposited in the Protein Data Bank (PDB) database under accession codes: XXX, XXX, XXX, and XXX. Corresponding cryo-EM maps, half maps, FSC curves and masks used have been deposited in the Electron Microscopy Data Base (EMDB) with accession codes: XXX, XXX, XXX, and XXX. The previously published structural models (PDB ids: 7WRG, 8UTN and 8UTS) were referenced in this work.

## Supplementary Information

**Supplementary Fig. 1.**
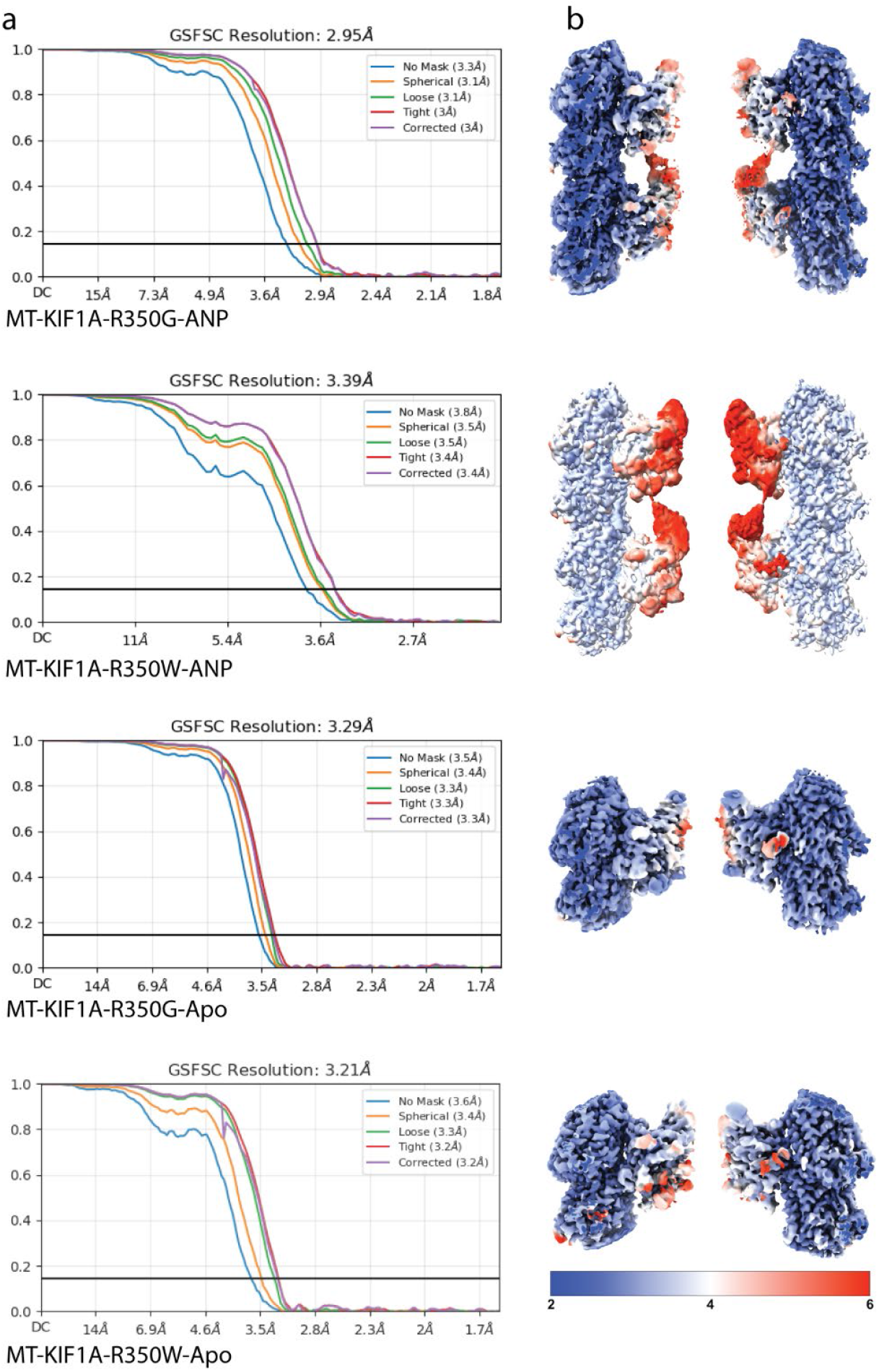
Fourier Shell Correlation (FSC) curves and local resolution. **a**. FSC curves. **b**. cryo-EM density map in surface representation colored according to local resolution. Color scale ranges from 2 Å (blue) to 4 Å (white) to 6 Å (red). From top to bottom FSC curves and local resolution maps correspond to : MT-KIF1A-R350G-ANP, MT-KIF1A-R350W-ANP, MT-KIF1A-R350G-apo and MT-KIF1A-R350W-apo.

**Supplementary Fig. 2.**
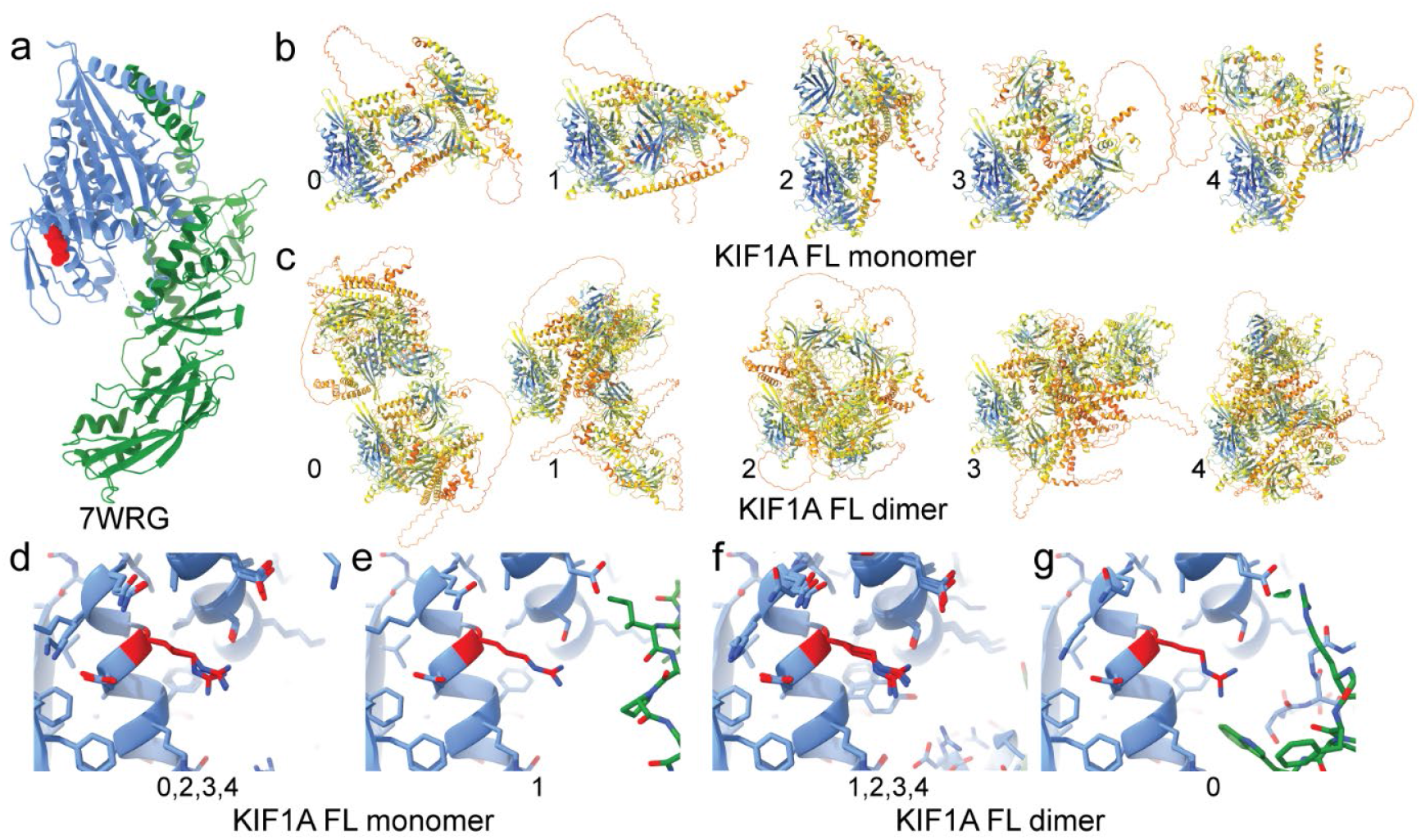
Kinesin-3 full-length (FL) models. **a**. *Ce*KLP6 crystal structure (PDB ID: 7WRG). The structure is shown in ribbon representation with the core motor domain in blue, the remainder of the molecule in green, and residue R354 (equivalent to human KIF1A R350) highlighted as red atom spheres. **b–c**. AlphaFold3-predicted models of human KIF1A in monomeric (b) and dimeric (c) forms. Structures are shown in ribbon representation and colored according to the AlphaFold estimated pLDDT confidence score (blue = high, red = low). **d–g**. Close-up views of the AlphaFold models, shown in ribbon representation with side chains depicted as stick atoms. The core motor domain is in blue, the remainder of the molecule in green, and residue R350 in red. Panels d and f show four overlapping models, aligned to the core motor domain, displaying similar configurations near residue R350, whereas panels e and g depict the more distinct models. Panels d–e correspond to monomer models, and panels f–g to dimer models. For clarity, only one motor head of each dimer model is shown, as both heads exhibit similar structures near residue R350. Notably, in none of the models does KIF1A residue R350 (or R354 in *Ce*KLP-6) contact other regions of the molecule.

**Supplementary Fig. 3.**
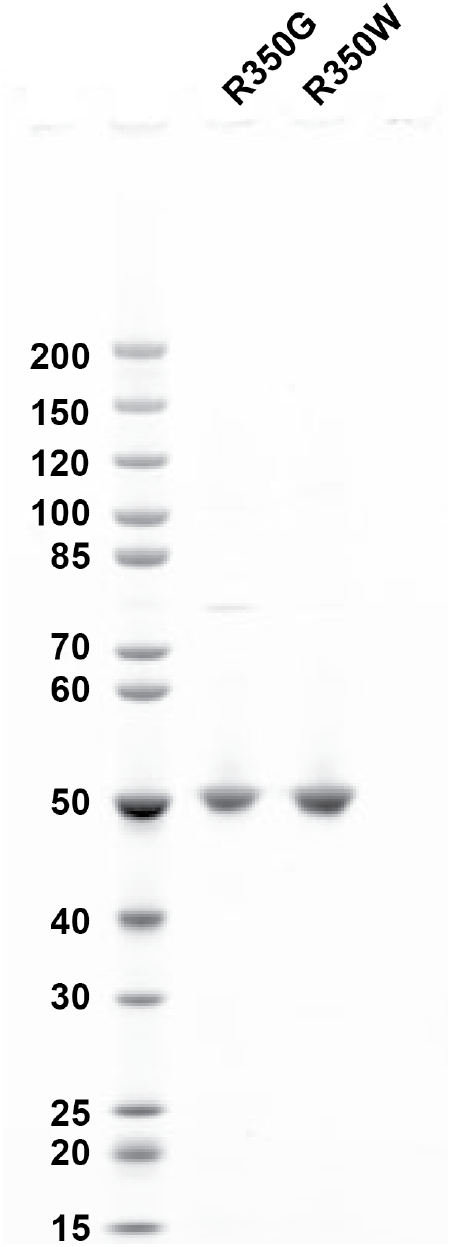
Polyacrylamide gel of KIF1A R350G and R350W.

